# Prioritizing peptides for targeted mass spectrometry experiments using deep learning

**DOI:** 10.64898/2026.05.21.727053

**Authors:** Shreyash Sonthalia, Bo Wen, Priank Dasgupta, Chris Hsu, Michael J. MacCoss, William Stafford Noble

## Abstract

One critical step in any targeted mass spectrometry experiment is selecting, from each protein of interest, a small number of peptides that respond well in the mass spectrometer and can serve as reliable proxies for protein quantification. Existing methods select target peptides either by relying on prior empirical measurements, limiting their applicability to previously observed peptides, or using machine learning to predict peptide behavior from sequence alone. However, current machine learning tools suffer from various limitations, including using detectability as an indirect proxy for intensity, relying on small training sets, or ignoring the precursor charge state. In this study, we introduce Bromo, a transformer-based deep learning model that ranks peptide precursors from a given protein by their relative response, taking charge state into account. Trained on millions of annotated peptide pairs derived from large-scale, publicly available data-independent acquisition mass spectrometry data, Bromo consistently outperforms existing sequence-based methods across diverse, independent datasets. Furthermore, we show that fine-tuning Bromo on experiment-specific data can account for differences in sample preparation, sample matrix, and instrument platform, all of which influence which peptides serve as optimal targets. This adaptability makes Bromo a practical tool for selecting target peptides for selected reaction monitoring and parallel reaction monitoring assay development across a wide range of experimental conditions.

## 1 Introduction

Targeted mass spectrometry (MS) coupled with liquid chromatography (LC) enables sensitive and reproducible quantification of proteins in complex biological samples by monitoring a selected set of target peptides using approaches such as selected reaction monitoring (SRM) ^1–3^ or parallel reaction monitoring (PRM) ^4,5^. Targeted mass spectrometry is widely used for applications such as biomarker detection, verification of results from untargeted data-dependent acquisition (DDA) and data-independent acquisition (DIA) analysis, and clinical assays, where quantitative precision and reproducibility are essential ^6–8^. One critical step in designing a targeted assay is selecting, from each protein of interest, a small number of peptides to monitor for the protein of interest. The quality of these peptide selections directly determines assay performance: ideal targets should yield strong, reproducible signals in the mass spectrometer and serve as reliable proxies for protein abundance.

A peptide’s response in the mass spectrometer depends on its amino acid sequence, the sample matrix, the instrument platform, and the acquisition method, making peptide selection a challenging problem. Early targeted assay development relied on DDA experiments to identify candidate peptides and to generate library spectra from which SRM transitions (i.e., pairs of corresponding precursor and product ions) could be derived ^9^. Although DDA’s stochastic sampling makes it unreliable as a direct predictor of peptide response in a targeted assay ^10^, the resulting library spectra provided a practical starting point for transition selection. To address the shortcomings of DDA-based selection, the field moved toward empirical refinement of assays directly in the target sample matrix using SRM or PRM ^11^, iteratively evaluating candidate peptides on the target instrument. However, this strategy becomes difficult when the target protein is near the instrument’s limit of detection, where it is often unclear whether the observed signal corresponds to the intended peptide of interest. To avoid this ambiguity, an alternative empirical strategy is to produce candidate proteins recombinantly via in vitro transcription and translation and then directly measure their tryptic peptides in isolation ^10,12^; this approach yields high-quality peptide selections but is time consuming and expensive. More recently, several groups have demonstrated that DIA data can efficiently inform peptide selection by providing comprehensive measurements of peptide behavior under conditions closely related to targeted acquisition, using DIA chromatogram libraries to select and schedule peptide targets for SRM and PRM assays ^7,13–15^. Although empirical approaches work well when relevant measurements are available, they cannot generalize to proteins that have not previously been observed and may not perform well in new experimental conditions.

An alternative strategy is to use machine learning to predict a peptide’s response directly from its amino acid sequence, removing the dependence on prior measurements. The earliest work in this direction used DDA data to predict which peptides will be reliably detected ^16,17^; however, as noted above, DDA-based peptide selection is suboptimal for targeted proteomics experiments. More recent tools, such as PeptideRanger ^18^, use random forest models trained on large, diverse collections of mass spectrometry experiments but still rely primarily on DDA data and predict detectability rather than quantitative intensity. Another related method, d::pPop, uses hand-crafted physicochemical peptide features together with a five-layer, fully connected neural network trained on DDA data to predict peptide observability scores and induce a within-protein ranking ^19^. One notable exception to the reliance on DDA is PREGO ^20^, which trains a neural network on features derived from a set of equimolar synthetic peptides measured by DIA; however, the small, synthetic training set limits the diversity of peptide behavior the model can capture. In general, existing sequence-based tools suffer from various limitations, including using detectability as an indirect proxy for response, relying on small training sets that may not capture the diversity of peptide behavior in biological samples, or ignoring the precursor charge state.

The approach we describe here addresses these limitations by framing peptide selection as a relative ranking task. To select peptides for a given protein, it is sufficient to rank the candidate peptides relative to one another rather than predict their absolute responses. Our deep learning model, Bromo, takes as input pairs of peptide precursors from the same protein and predicts which will yield a larger quantitative response in a DIA experiment. This pairwise ranking model can then be used to induce a full ranking of peptides for a given protein. Bromo uses a transformer architecture ^21^ and was trained on millions of peptide pairs derived from large-scale, publicly available DIA data. We evaluate Bromo against existing sequence-based methods across diverse, independent DIA datasets, and assess whether fine-tuning can adapt the model to new experimental conditions.

## 2 Methods

### 2.1 The Bromo deep learning model

Bromo is a deep learning model that takes as input pairs of peptide precursors (peptide sequences and associated charge states) from the same protein and predicts whether the first precursor will yield a larger quantitative response than the second precursor in a DIA mass spectrometry experiment. The model consists of twin transformer encoders, sharing the same architecture and parameters, followed by fully-connected layers for binary classification. The encoder input is a list of integers, where each integer represents the token for an amino acid. This input is extended with a pad token to a maximum peptide length of 30.

To arrive at the architectural and optimization hyperparameters, we performed a joint hyperparameter search with Optuna (version 4.8.0) ^22^. Each trial samples a candidate configuration over the transformer embedding dimension, number of attention heads, number of Transformer layers, feed-forward dimension, charge-embedding dimension, classification-head hidden dimension, dropout rate, learning rate, weight decay, and label-smoothing coefficient, over the search ranges in Table 1. Trials that sample an incompatible embedding dimension/attention-head combination (i.e., where the embedding dimension is not divisible by the number of heads) are pruned. For each remaining trial, a model is trained with the sampled configuration for two epochs as a fast proxy for full training and scored by its validation AUROC. Trials are proposed with Optuna’s Tree-structured Parzen Estimator (TPE) sampler, and the configuration from the best of 10 trials is carried forward to the final training run. This search yielded the architecture and optimization settings used for the final Bromo model described below.

**Table 1:** Optuna search space for architectural and optimization hyperparameters.

| Hyperparameter | Type | Search range |
| --- | --- | --- |
| Learning rate | log-uniform | $[10^{-5}, 10^{-3}]$ |
| Weight decay | log-uniform | $[10^{-4}, 10^{-1}]$ |
| Dropout rate | uniform | $[0.1, 0.5]$ |
| Label smoothing | uniform | $[0.0, 0.2]$ |
| Embedding dimension ($d_{\text{model}}$) | categorical | $\{64, 128, 256\}$ |
| Attention heads | categorical | $\{4, 8\}$ |
| Transformer layers | integer | $[2, 6]$ |
| Feed-forward dimension | categorical | $\{256, 512, 1024\}$ |
| Charge-embedding dimension | categorical | $\{4, 8, 16\}$ |
| Classifier hidden dimension | categorical | $\{128, 256, 512\}$ |

The encoder consists of four layers: an embedding layer with output dimension 256, a positional encoder layer for each token, and four transformer layers, each with 8 attention heads and a feedforward dimension of 512. The output token representations are aggregated into a single 256-dimensional vector via learned attention pooling, where a learned scoring function assigns a weight to each token position and the final representation is the weighted sum. The vector representations of the two peptides are concatenated together with the two charge state embeddings (dimension 16), yielding a 544-dimensional joint representation, which is fed into two fully connected layers with dimensions 128 and 2, respectively. Dropout (rate of 0.156) was applied to the transformer layers and the fully connected classifier layers. The model outputs logits over two classes, corresponding to which of the two peptide-charge combinations is predicted to yield a larger MS2 intensity value. The model optimizes the cross-entropy loss function using the AdamW optimizer ^23^. After training, the best-performing model was selected using the area under the ROC curve, calculated on the validation set using the human-astral dataset.

The final model is trained on a single NVIDIA L40 GPU for 15 epochs with the AdamW optimizer using the tuned learning rate, weight decay, and label-smoothing coefficient, and a batch size of 4096. Gradient norms are clipped to a maximum of 1.0, and the learning rate is reduced by a factor of 0.5 (`ReduceLROnPlateau`) whenever the validation loss fails to improve for two consecutive epochs. The model is evaluated on the held-out validation set every 100 training steps, and the checkpoint with the highest validation AUROC is retained as the final model. The training time is approximately 45 hours. The corresponding runtime for applying the model to a new dataset or protein list is shown in Supplementary Figure 3.

For reproducibility, we fix a single random seed (default 42, exposed via `--seed`) that is used to seed PyTorch’s CPU and CUDA random number generators (`torch.manual_seed`, `torch.cuda.manual_seed_-all`) before model initialization and again before each Optuna trial. NumPy’s random number generator is used for protein-level subsampling, Optuna’s TPE sampler is used during hyperparameter search, and the `random_state`of all pandas-based sampling operations is used to construct the training data. We disable cuDNN’s non-deterministic autotuner (`torch.backends.cudnn.benchmark=False`) and enable de-terministic algorithm selection (`torch.backends.cudnn.deterministic=True`) so that GPU computations are reproducible given a fixed seed.

### 2.2 Deriving precursor-level and peptide-level rankings from precursor-pair binary classification

The goal of Bromo is to induce a ranking over all the peptides (in each possible charge state) in the protein. To do this, binary classification results for each precursor pair within a protein are used to derive a precursor-level ranking. We induce a ranking by running inference for each peptide+charge pair combination in the protein and sorting by the number of times they are assigned a positive label by the model, with ties broken by the average predicted score assigned to the peptide. For each pair, inference is run for both directions (i.e., (*A, B*) and (*B, A*)) and an equally weighted convex combination of the two predictions is used as the representative score for the pair. The tie-breaking step is necessary because the model does not always obey transitivity.

For comparison to existing methods that do not consider charge state^18,20^, we derive a peptide-level ranking by choosing a representative precursor for use in peptide-level ranking. We do this by taking the precursor-level ranking, keeping only the precursor with the highest true ranking as the representative peptide, and then sorting by the average predicted score assigned to the peptide to break ties.

### 2.3 Datasets

A total of seven datasets were used in the study (Table 2). The first dataset, human-astral, was used for pretraining because this dataset contained the largest number of detected peptides. We used the other six datasets for test-sets and/or finetuning. The yeast datasets were also from the same study as human-astral and were used for finetuning and testing the base Bromo model because they contained protein groups and peptides not found in the human datasets (ProteomeXchange: PXD056793) ^24^. The human-pan dataset was generated from 949 cancer cell lines across 28 tissue types analyzed by SCIEX 6600 TripleTOF mass spectrometers. This dataset includes a total of 6864 DIA runs, and was also used for fine-tuning the model because this dataset contained the largest number of independent DIA runs, and thus precursor-pair labels with lowest variance. DIA-NN peptide detection results were downloaded from PRIDE with identifier PXD030304.

**Table 2:** Summary of datasets used in the study. A peptide is counted in the “Peptides” column if it participates in at least one peptide pair. Precursor pairs are defined as all unique pairs of precursors from the same protein, prior to data augmentation to induce equivariance to the order of the input peptides.

| Dataset | Instrument | Sample | Runs | Protein groups | Precursors | Precursor pairs |
| --- | --- | --- | --- | --- | --- | --- |
| human-astral | Orbitrap Astral | HeLa cells | 4 | 8070 | 443778 | 18384919 |
| human-pan | 6600 TripleTOF | Cancer cell lines | 6864 | 6777 | 280765 | 10504865 |
| yeast-astral | Orbitrap Astral | Yeast | 4 | 4710 | 317958 | 12777431 |
| human-lumos | Orbitrap Fusion Lumos | HeLa cells | 3 | 5879 | 230799 | 6892272 |
| yeast-lumos | Orbitrap Fusion Lumos | Yeast | 3 | 4100 | 187053 | 5346828 |
| human-exploris | Exploris 480 | HeLa cells | 4 | 5521 | 220125 | 6964014 |
| yeast-exploris | Exploris 480 | Yeast | 4 | 3971 | 185481 | 5679852 |

The two other human datasets were generated from different instruments and were also from the same study as human-astral (ProteomeXchange: PXD056793) ^24^. These datasets were also used for fine-tuning. Each dataset was analyzed with DIA-NN (version 1.8.1) ^25^, using the same protein database and search settings as the original study ^24^. Only precursors and proteins with a DIA-NN q-value *≤* 0.01 were retained for analysis. Throughout, we used the normalized precursor intensity (“Precursor.Normalised”) from DIA-NN’s main report as the precursor intensity.

### 2.4 Assigning labels to peptide pairs

For a given set of one or more MS runs, a collection of labeled precursor pairs was created in three steps. For each individual MS run and protein group, we first generate all possible precursor pairs, excluding self comparisons and excluding any precursor that maps to multiple proteins. This is done by performing *in silico* tryptic digestion of each protein sequence using trypsin without allowing missed cleavages, peptide length 7–30, and assigning charge states of 2, 3, and 4 to each peptide. Note that this restriction applies only to the *in silico* digestion: peptides detected by DIA-NN are retained even if they contain a missed cleavage. Second, for each pair of peptide-charges (*A, B*), if both *A* and *B* were detected in the run, then a label is assigned based on whether *A* has a higher observed intensity (label “1”) or a lower observed intensity (label “0”). If *A* was detected but not *B*, then the label “1” is assigned, and vice versa if *B* was detected but not *A*. If neither *A* nor *B* was detected in the run, then the pair is discarded. As a form of data augmentation to induce equivariance to the order of the input peptides, each pair is included in the dataset twice (i.e., (*A, B*) and (*B, A*)), with opposite labels.

Finally, a consensus label is assigned to each pair across all the runs in the dataset. Say that, after the two-step procedure outlined above, a given pair (*A, B*) has a label assigned for *n* runs, and that the label is positive in *m* of those runs. For large datasets (human-pan in Table 2), we use the binomial distribution to calculate the probability of observing at least *m* heads in *n* flips of a fair coin:

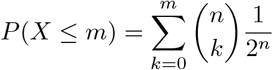

We set the consensus label to “1” if 1 *−P* (*X ≤ m −*1) *< τ* and “0” if *P* (*X ≤ m*) *< τ*. In this work, we use *τ* = 0.05. The binomial distribution method was only used for peptide pairs that were detected in at least five runs, with a simple majority voting scheme used for the remaining peptide pairs. For the remaining, smaller datasets, we assign the consensus label using the majority voting scheme.

### 2.5 Splitting the data for training, validation, and testing

For the human-astral dataset, we randomly split the protein groups into training, validation, and testing sets with a ratio of 0.6, 0.2, and 0.2, respectively. The training set was used to train the model, the validation set was used to decide upon training convergence, and the testing set was used to evaluate the model performance.

For other human datasets used for finetuning, we created train/validation/test splits that respect the original splits created for the human-astral dataset. This is done by identifying any protein groups and then peptide pairs that are shared between the new dataset and the human-astral training set. We then split the remaining protein groups randomly, to achieve overall train/validation/test ratios of 0.6, 0.2, and 0.2. For the yeast datasets used for finetuning, we created train/validation/test splits on protein groups with a ratio of 0.6, 0.2, and 0.2, respectively.

### 2.6 Model fine-tuning

Fine-tuning adapts a pretrained Bromo model to a new experimental context by continuing training on peptide pairs drawn from a target dataset generated from the new context. We initialize the model with the weights of the model pretrained on the human-astral dataset, using the checkpoint that achieved the highest validation AUROC. The architecture is unchanged, and all model parameters are updated. Fine-tuning uses the train, validation, and test splits described above, so that the protein groups used for fine-tuning are disjoint from those used for evaluation and respect the splits established for the human-astral dataset.

Fine-tuning optimizes the same cross-entropy objective, with a label smoothing of 0.058, using the same AdamW optimizer as pretraining. We reuse the learning rate (3.8 *×*10^*−*5^) and weight decay (6.8 *×*10^*−*3^) and a batch size of 4096, and we fine-tune for five epochs rather than the 15 used during pretraining. All of these hyperparameters are the same as the ones selected during hyperparameter tuning during pretraining. Gradient norms are clipped to a maximum norm of 1.0, and the learning rate is reduced by a factor of 0.5 whenever the validation loss fails to improve for two consecutive epochs. We evaluate the model on the validation set every 100 training steps and retain the checkpoint with the highest validation AUROC as the final fine-tuned model.

### 2.7 Performance measures

To evaluate the performance of the various models, we designed a performance measure that quantifies the quality of a given set of selected peptides. Based on the ranking of peptides produced by a given method, we select a specified number *k* of top-ranked peptides to target, discarding any protein group with 10 or fewer unique peptides. We then compare the top-*k* predicted peptides to the gold standard ranking of peptides by their observed intensities. The performance measure, the top-*k* accuracy (TKA), is the fraction of predicted peptides that are included in the top *k* according to the gold standard:

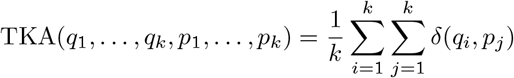

where *q*_1_, …, *q*_*k*_ are the top *k* predicted peptides, *p*_1_, …, *p*_*k*_ are the top *k* peptides in the gold standard, and *δ*(*q, p*) is 1 if *q* = *p* and 0 otherwise. The value of TKA ranges between 0 and 1, with larger values being better. In practice, we compute average TKA values across all proteins in a given dataset, and we repeat the calculation for varying values of *k*. To assess the statistical significance of a given observed difference in performance between two methods, we use a paired t-test on the per-protein differences in TKA.

### 2.8 Baseline methods

We compared the Bromo model to four baseline methods: a chance-level baseline, PREGO, PeptideRanger and XGBoost. All four baseline methods were applied to the same set of peptides analyzed by Bromo, including peptides up to length 30.

For the top-*k* accuracy (TKA) metric, we define chance-level performance analytically rather than by fitting a null model. For a protein with *N* distinct peptides passing the same filtering criteria used to compute TKA (at least 10 unique peptides and at least 7 detected peptides), a uniformly random ranking of the *N* peptides is expected to place *k/N* of the true top-*k* peptides within its own top-*k* list, since two independent random *k*-subsets of an *N* -element set have an expected fractional overlap of *k/N*. We use this quantity, TKA_chance_(*k*) = *k/N*, as the chance-level curve for each protein, and we aggregate it across proteins in the same way as the method curves (mean over proteins at each *k*), yielding a single reference curve that is plotted alongside bromo and the baseline methods (dashed gray line, e.g., Figure 2).

PREGO was downloaded from https://github.com/briansearle/intensity_predictor. We used PREGO’s `classify`command to generate a score *s*(*p*) for each peptide, which can be used to rank peptides within a protein. where *s*(*p*) is the score assigned by PREGO to peptide *p*.

PeptideRanger was downloaded from https://github.com/rr-2/PeptideRanger. The function `create_peptidome`was first used to generate a list of peptides from each protein, using the following parameters: `missed cleavages = c(0,0)`, `synth_peps = FALSE`and `aa_range = c(7,30)`. Then the function `prioritize_peptides`was used to predict the detectability of each peptide with the following parameters: `max_n = 20`, `prediction_model = RFmodel_ProteomicsDB`, `priorities = “RF_score”`and `priority_thresholds= 0`. The detectability score (`RF_score`) was then used to rank pairs of peptides from the same protein.

As a baseline, we trained an XGBoost model on the same peptide-pair classification task as Bromo. Rather than learned embeddings, each precursor is represented by its unigram amino-acid composition (counts over the 21-letter amino-acid alphabet) together with one-hot indicators for its N-terminal and C-terminal residue. The composition and terminal features of the two peptides are concatenated with their two (unembedded) charge states into a single 128-dimensional feature vector per pair, and the pair label is taken directly from the input table. XGBoost hyperparameters were tuned with Optuna, using validation AUROC—computed by scoring each trial’s fitted booster on a held-out validation set—as the objective. The search covered maximum tree depth ([3, 10], integer), learning rate ([10^*−*3^, 0.3], log-uniform), subsample and column-subsample fractions ([0.5, 1.0], uniform), minimum child weight ([1, 10], integer), *L*_1_ and *L*_2_ regularization strength ([10^*−*2^, 10], log-uniform), and the number of boosting rounds ([50, 500], integer), with the binary cross-entropy objective and a fixed random seed held constant across trials. Trials were proposed with Optuna’s Tree-structured Parzen Estimator sampler across 10 trials; no intermediate pruning was used. The hyperparameters and boosting-round count from the best trial were saved and used to refit the final model on the full training set, with the validation set used only for hyperparameter selection and not included in the final fit.

### 2.9 Peptide ranking comparison across experimental replicates and instruments

To evaluate precursor ranking consistency across experimental replicates and instrument platforms, we compared precursor rankings derived from pairs of DIA runs acquired on Orbitrap Astral, Orbitrap Fusion Lumos, Exploris 480, and SCIEX 6600 TripleTOF instruments. For each comparison, we restricted the analysis to proteins for which the same two peptides were detected in both runs and generated all within-protein precursor pairs. One run was treated as the observed ranking, with pairwise labels assigned according to which precursor had the larger DIA-NN normalized precursor intensity, and the other run was treated as the predicted ranking, with pairwise scores defined by the corresponding precursor intensity ratios. For comparisons across experimental replicates, the two runs were drawn from the same dataset. For cross-instrument comparisons, the observed ranking was derived from DIA data acquired on one instrument and the predicted ranking was derived from DIA data acquired on a different instrument. We then quantified agreement between the observed and predicted rankings using the same pairwise ranking analysis used throughout the manuscript.

## 3 Results

### 3.1 The Bromo model learns to rank peptide pairs

To effectively prioritize peptides within a protein for detection in a targeted proteomics experiment, we built a deep learning model called Bromo that can be used to rank peptide precursors (i.e., peptide sequences with specific charge states) within a protein based on their predicted MS2 response signals in DIA experiments. As shown in Figure 1a, Bromo takes as input an ordered pair of precursors from the same protein and predicts whether the first precursor will yield higher intensity than the second precursor in a DIA experiment. The underlying assumption is that peptides from the same protein are expected to have the same concentration in the sample, and the differences in their response signals are mainly due to the sequence features of the peptides and the variation introduced during sample preparation and peptide ionization. Note that this assumption concerns the abundances of the peptides, not the intensities they ultimately produce: sample preparation effects such as incomplete peptide cleavage, incomplete alkylation, and oxidation are peptide-specific response differences of the kind the model is trained to predict. In general, for a set of peptides derived from the same protein, selecting peptides with higher intensities in a DIA experiment is preferred for targeted proteomics experiments ^13,14,20^. We hypothesize that a model trained directly on peptide response signals can rank peptides for targeted proteomics experiments more effectively than models that predict simpler observability metrics such as peptide detectability ^17,18^. In addition, our approach does not rely on synthetic datasets for model training; instead, it can be trained directly on DIA datasets from complex samples (Figure 1b). Bromo can also be fine-tuned on data generated from different MS instruments to improve performance under specific experimental settings.

**Figure 1:**
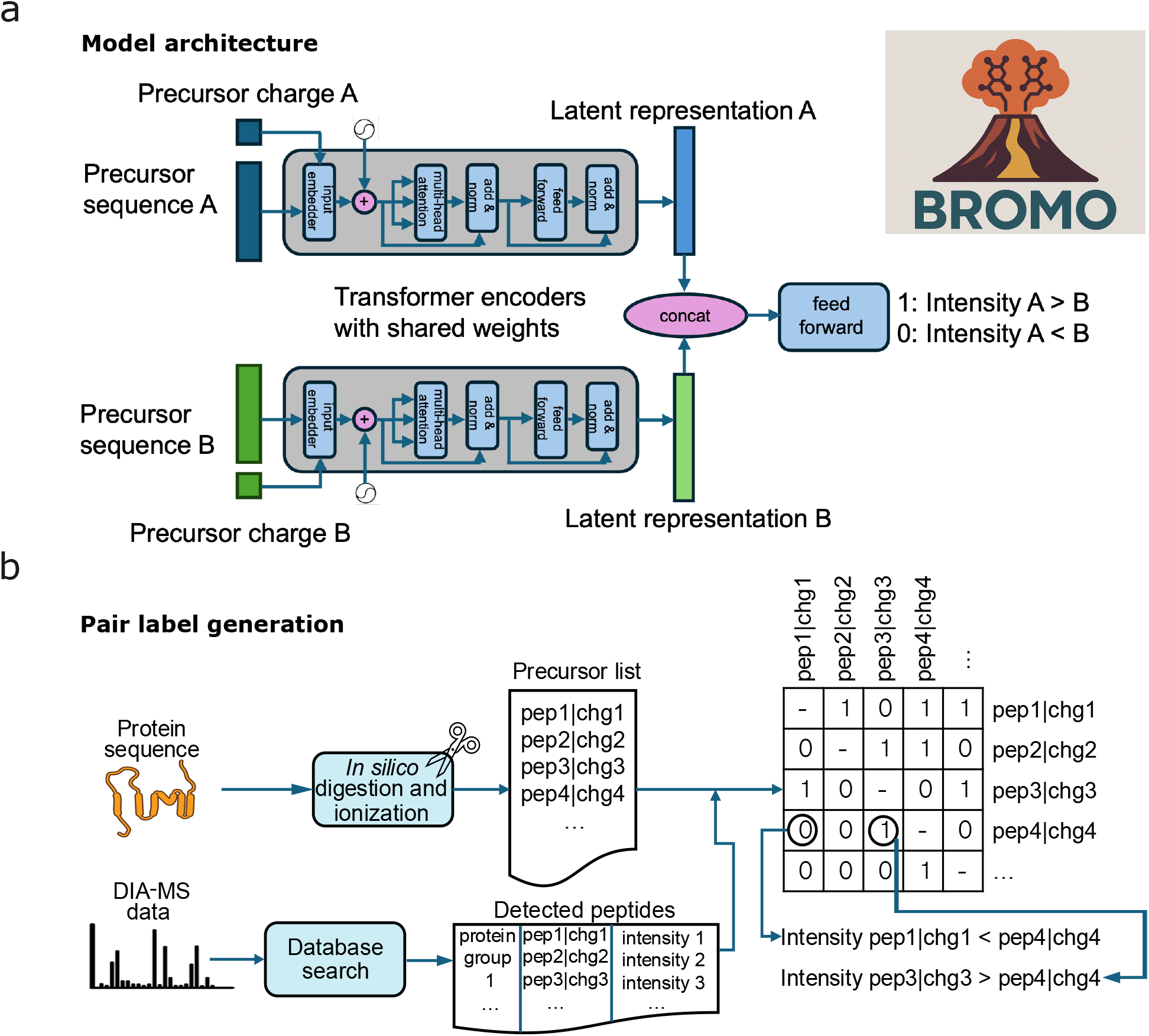
Overview of peptide ranking. (a) Schematic overview of the Bromo model architecture. (b) Training data generation for peptide ranking. The binary label at position (*i, j*) in the label matrix indicates whether precursor *i* exhibits higher intensity than precursor *j*.

### 3.2 Bromo outperforms existing methods for peptide ranking

To evaluate the performance of our approach, we benchmarked Bromo across three independent datasets acquired on three different MS platforms (Orbitrap Astral, Orbitrap Fusion Lumos, and Orbitrap Exploris 480), as shown in Table 2. We pretrained Bromo on proteins detected in a human DIA dataset (human-astral) and tested its performance on yeast DIA datasets generated under identical LC-MS/MS experimental conditions (yeast-astral, yeast-lumos, and yeast-exploris). This evaluation setup ensures that high performance on the test set reflects genuine generalization rather than memorization of training peptides. We first compared Bromo against a baseline method that uses a standard machine learning classifier, XGBoost ^26^ (see Methods for details), trained and evaluated using the same data splits. Bromo consistently outperformed the XGBoost baseline, achieving significantly higher top-*k* accuracy (TKA) across all three evaluation sets (Figure 2a–c). Because XGBoost operates on a fixed compositional encoding while Bromo learns directly from the peptide sequence, this comparison reflects both the model architecture and the input representation, and is not intended to isolate the contribution of the transformer architecture.

**Figure 2:**
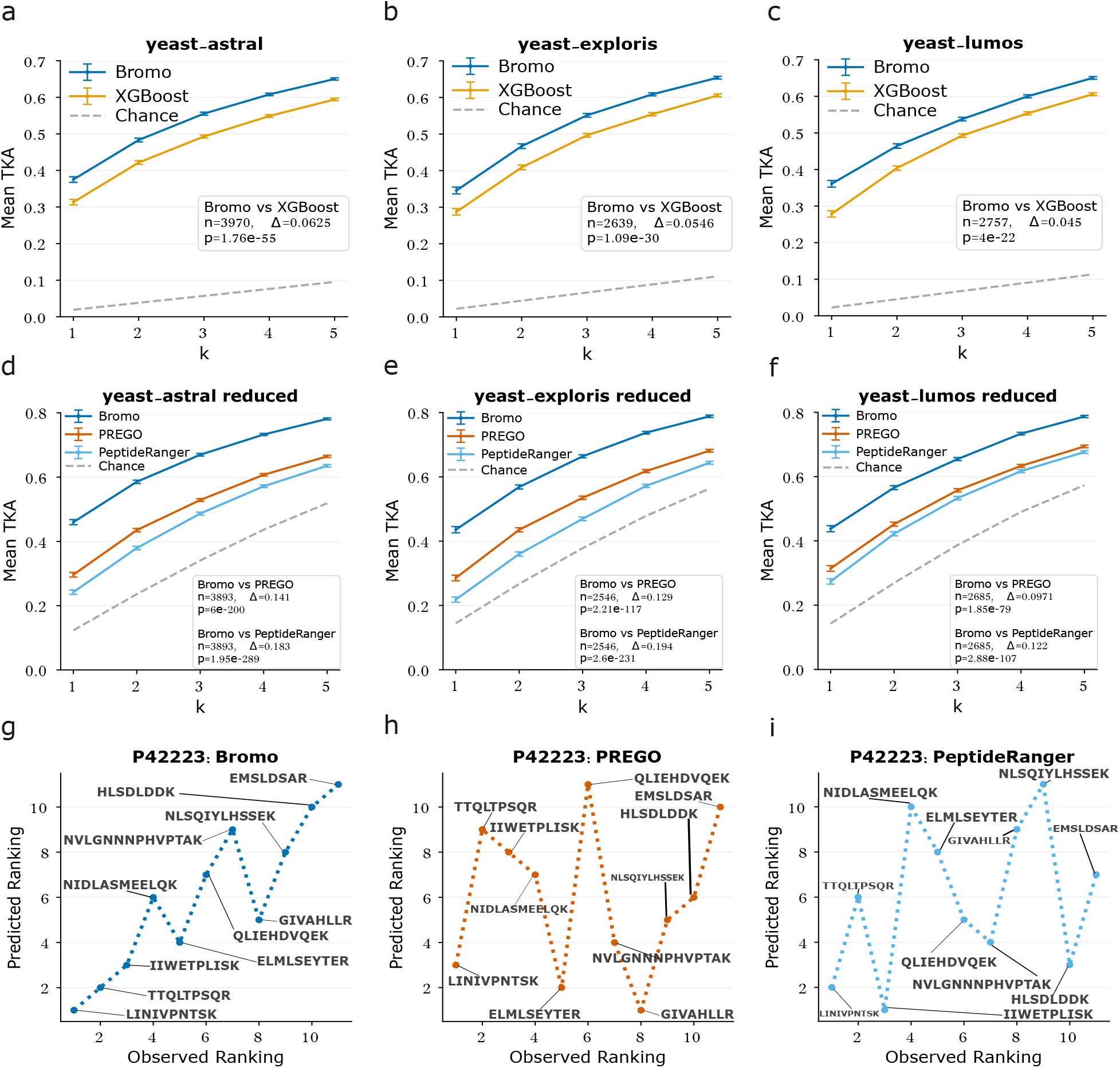
Peptide ranking performance on three DIA datasets. In panels **(a)–(f)** The figure plots the top-*k* accuracy (TKA) for *k* = [1, 5] on paired yeast datasets from three different MS instruments (Astral: **(a)** and **(d)**, Exploris: **(b)** and **(e)** and Lumos: **(c)** and **(f)**). The blue curves always correspond to Bromo, orange curves correspond to the baseline XGBoost model, red curves correspond to PREGO, and light blue curves correspond to PeptideRanger. Dashed gray lines correspond to the chance-level baseline. In panels **(d)–(f)**, datasets are reduced by only comparing peptides output by the PREGO and PeptideRanger tools. **(g)–(i)** Rankings of peptides from protein P42223 using Bromo, PREGO, and PeptideRanger, respectively.

We further examined how ranking performance varies with the number of candidate tryptic peptides in a protein group. Stratifying the mean TKA at *k* = 3 by the number of candidate peptides up to the 95th percentile of the distribution (Supplementary Figure 1), we found that performance was highest for protein groups with few candidate peptides and declined as the number of candidates increased, consistent with the expectation that ranking becomes more difficult—and chance-level TKA lower—as the number of candidates grows. Across this range, Bromo generally maintained an advantage over the XGBoost baseline, with the difference most pronounced for protein groups containing larger numbers of candidate peptides in the Exploris and Lumos datasets.

We next benchmarked Bromo against PREGO and PeptideRanger across the same three evaluation sets. Because these tools do not natively model precursor charge states, we derived peptide-level rankings from Bromo’s precursor-level predictions for a fair comparison (see Methods). Comparisons were further restricted to peptides present in the rankings output by each tool. On this reduced yeast-astral dataset, Bromo yielded significantly higher TKA values at *k* = 3 than both PREGO (paired *t*-test, *p* = 6*×*10^*−*200^)and PeptideRanger (paired *t*-test, *p* = 1.95 *×* 10^*−*289^)(Figure 2d–f). An illustrative example of peptide ranking by the three methods is shown in Figure 2g–i, in which a randomly sampled protein group from the yeast-astral dataset was sampled and peptide-level rankings were computed.

Notably, while PREGO was explicitly designed to model peptide ranking, PeptideRanger was trained to predict general peptide observability (i.e., whether a peptide is detected across a broad array of mass spectrometry experiments) and uses this detectability as a proxy for ranking. This fundamental difference in training objectives likely accounts for PeptideRanger’s relatively poor performance on this targeted selection task.

### 3.3 Instrument-specific fine-tuning mitigates cross-platform variability

Mass spectrometry instrument characteristics can profoundly influence peptide detectability and relative response. To quantify this variability and evaluate strategies to overcome it, we assessed the consistency of peptide rankings across different MS platforms and tested whether instrument-specific fine-tuning improves Bromo’s predictive performance.

We first established baselines for intra-instrument and inter-instrument consistency of peptide rankings. To do this, we began by comparing rankings derived from technical replicate measurements on a single instrument platform. We arbitrarily selected one replicate as the gold standard and computed TKA for the ranking from the other replicate relative to that gold standard. As shown in Figure 3a–d, TKA values at *k* = 3 on the four datasets, each derived from a distinct MS platform, ranged between 0.850 and 0.930. This result confirms that peptide ranking remains highly consistent across independent runs acquired with identical instrument settings. Next, we repeated this experiment using replicates acquired on different types of instruments. Again, we selected one instrument platform as the gold standard and computed TKA based on the ranking from the other instrument platform. This inter-instrument analysis revealed a substantial reduction in ranking consistency, measured using TKA at *k* = 3, to a range of 0.528 to 0.745 (Figure 3a–d), highlighting the considerable variability in rankings produced by different mass spectrometer platforms.

**Figure 3:**
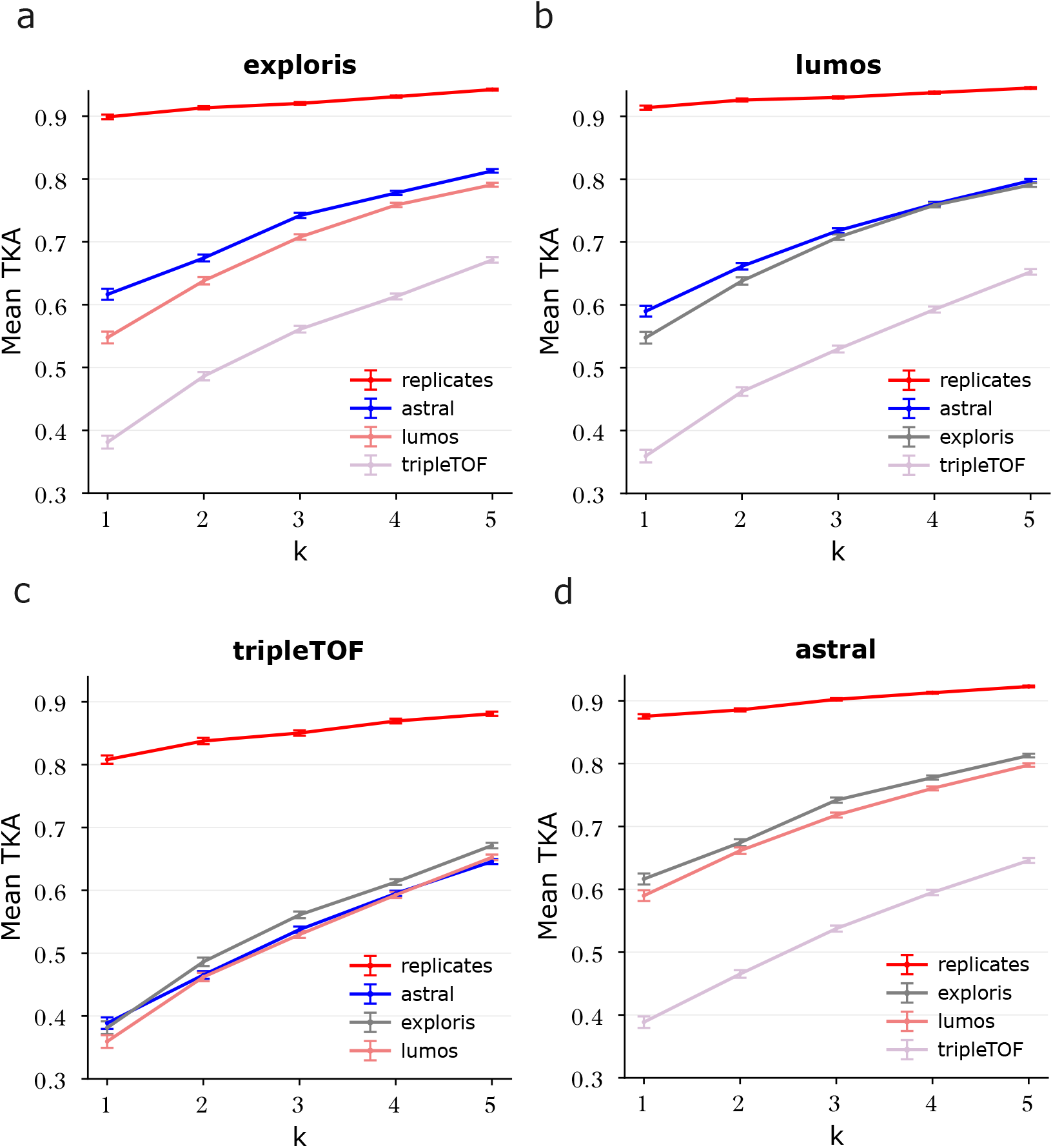
Peptide response varies across different instruments. **(a)–(d)** Peptide ranking consistency across replicates and instruments. All data used in these analyses were from human cell line samples. For intra-instrument comparisons, shown as red lines, one replicate from each dataset was taken as the gold standard and the other replicate from the same dataset was taken as the test ranking. For cross-instrument comparisons, the instrument platform listed at the top of each plot is used as the gold standard, and TKA is calculated with respect to replicates from three different platforms, listed in the legend of each plot.

We next investigated whether fine-tuning could adapt a pretrained Bromo model to a different type of instrument. To this end, we compared the performance of a Bromo model trained using the human-astral dataset with the performance of the same model after it had been fine-tuned using data from the same instrument as the test data (Exploris, Lumos, or SCIEX TripleTOF). We observed that, in each case, performance after fine-tuning was either better than or not significantly different from that of the pretrained model, though the degree of improvement varied (Figure 4). For the human-pan dataset, generated on a SCIEX TripleTOF, the TKA at *k* = 3 improved by 0.0835 (from 0.501 to 0.585). We note that this improvement is statistically significant (paired *t*-test, *p* = 3.4 *×*10^*−*21^). Significant improvements were also observed for the human-exploris and human-lumos datasets (0.0247 and 0.0255 respectively), but improvements were relatively small.

**Figure 4:**
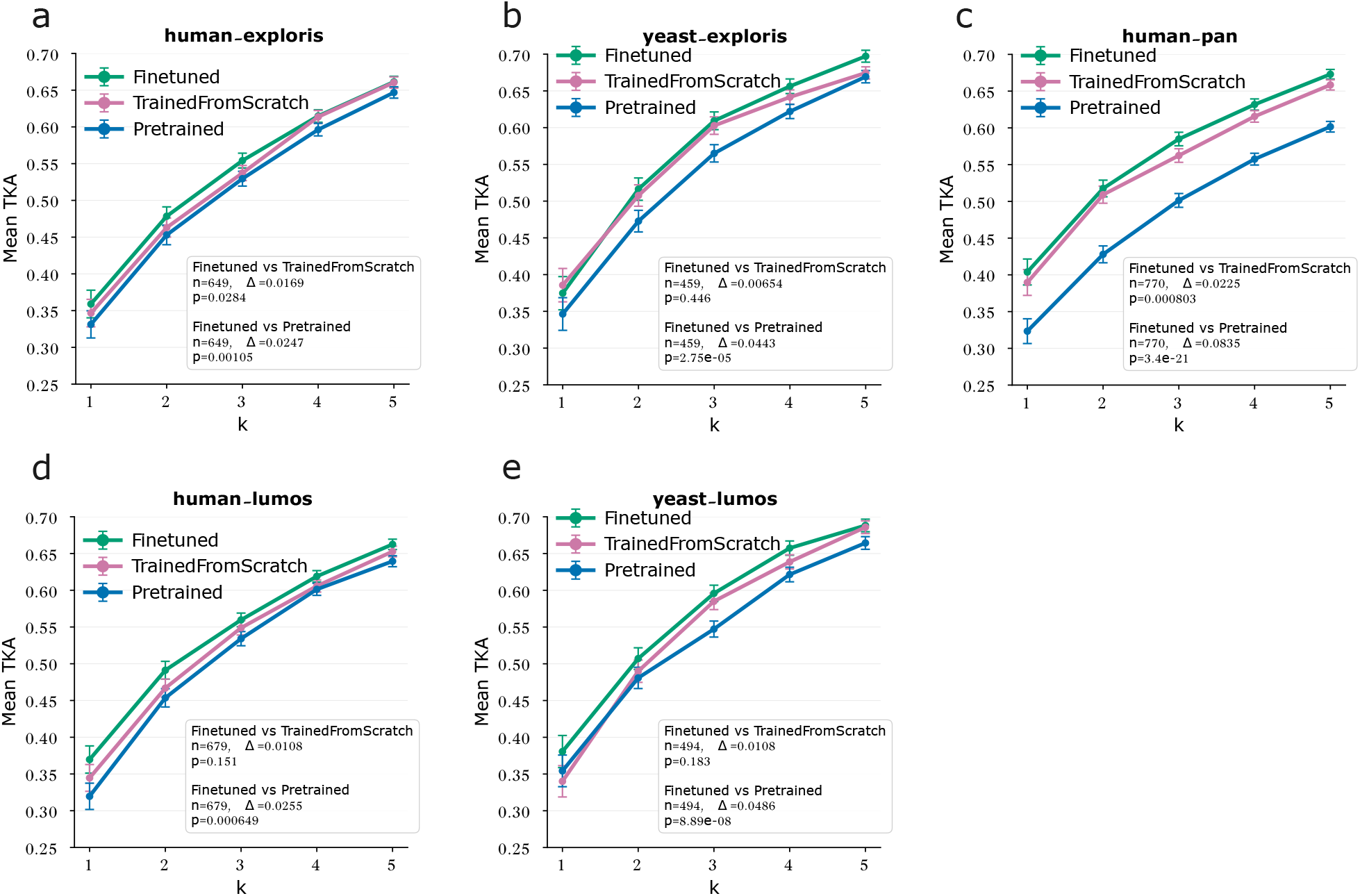
Evaluating model fine-tuning for peptide ranking on different datasets. Three types of models (fine-tuned models, models trained from scratch, and a pretrained model) were compared on five datasets: **(a)** human-exploris, **(b)** yeast-exploris, **(c)** human-pan, **(d)** human-lumos, and **(e)** yeast-lumos.

We then compared fine-tuned models with models trained from scratch on each dataset. As shown in Figure 4, fine-tuning yielded better or comparable performance on most datasets relative to models trained from scratch. Notably, while the human-pan and human-exploris datasets benefited significantly from fine-tuning, the Lumos human dataset did not exhibit similar improvements. Fine-tuning, however, was never worse than the pretrained model or the model trained from scratch on each dataset.This result suggests that while fine-tuning is a viable strategy for mitigating cross-platform variability in some contexts, its overall efficacy depends on the specific instrument and dataset characteristics.

### 3.4 Bromo’s performance scales with increasing training data

To evaluate the impact of training set size on performance, we plotted ranking accuracy as a function of the number of protein groups used for training. This analysis, performed on both the human-astral and human-pan datasets, allowed us to compare how well each model scales with additional data. As illustrated in Figure 5, Bromo’s performance exhibits steady improvement as the number of training peptide pairs increases. In contrast, the performance of the XGBoost baseline model quickly plateaus. This divergence highlights the capacity of Bromo’s deep learning architecture to effectively leverage increasing amounts of training data for superior peptide ranking, an advantage that traditional machine learning methods fail to capture.

**Figure 5:**
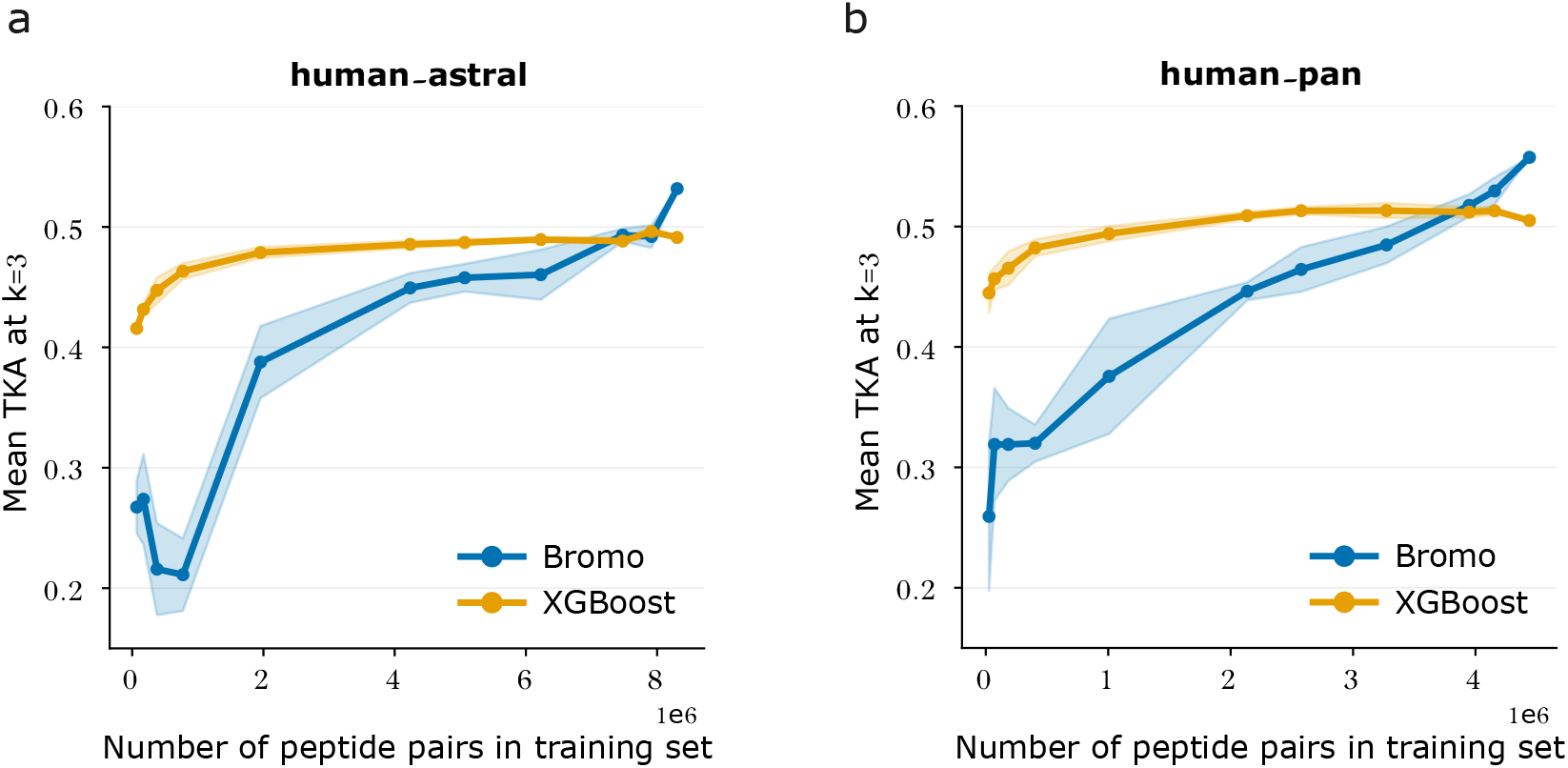
Evaluating the impact of training data size on model performance. Mean TKA at k = 3 relative to peptide pair training set size was assessed for Bromo and XGBoost for (a) the human-astral dataset and (b) the human-pan dataset. Shaded area demonstrates standard deviation across 5 random subsamples of the training set.

### 3.5 Comparing binomial and majority voting labeling schemes

Bromo is trained on peptide precursor pairs with binary labels indicating which precursor yields a larger quantitative response. Because a given pair may be observed in multiple MS runs, and the observed relative responses may vary across runs, we use either majority voting or a binomial procedure to assign a single consensus label across runs. In general, a desirable property of a labeling scheme is that, for a given pair of peptides, the assigned label should be unlikely to change as the number of observations increases. To quantify the consistency of each of our two labeling schemes with respect to the number of observations, we carried out a downsampling simulation. Using the human-pan dataset, we assigned each peptide pair a “full data” label based on all 6864 available runs, using either the binomial assignment or the majority voting procedure depending on the number of runs in which the peptide pair was detected. We then randomly selected 20 runs, re-computed the labels, and counted how many peptide pairs received a “downsampled” label that differed from the “full data” label; we repeated this procedure for 2, 3, 4, 5, 6, 7, 8, 9, 10, 12, 15, 17 runs. The resulting curves (Supplementary Figure 2) suggest that while the binomial distribution labeling scheme yields labels that are more consistent, the majority voting scheme does not perform significantly worse: for both schemes, over 98% of peptide pairs detected in at least 5 runs received the same label across all 20 runs.

## 4 Discussion

Bromo addresses a central problem in targeted proteomics assay development: selecting, from the candidate peptides for a protein of interest, those most likely to produce strong and reliable mass spectrometry signals. Rather than predicting peptide response signals on an absolute scale, Bromo directly models the relative response of peptide precursors (i.e., peptide sequences with specific charge states) from the same protein. This formulation is well matched to the practical decision made during assay design, where the goal is usually not to estimate an absolute peptide intensity but to choose a small number of high-responding peptides. Across independent DIA datasets generated on multiple instrument platforms, this pairwise ranking approach outperformed both a conventional XGBoost baseline trained on the same data and the existing sequence-based tools PREGO ^20^ and PeptideRanger ^18^. These results suggest that DIA datasets with deep peptide and protein coverage provide a useful empirical basis for learning peptide response properties relevant to targeted assay development.

Three design choices drive this performance. First, Bromo uses observed precursor intensity, rather than peptide detectability, as the training signal. Detectability is useful for identifying peptides that are likely to be observed, but it does not distinguish among the subset of detectable peptides that differ substantially in quantitative response; this distinction is reflected in Bromo’s large advantage over PeptideRanger. Second, Bromo models peptide precursors, including charge state, rather than peptide sequences alone, because different charge states of the same peptide can have different ionization efficiencies, fragmentation behaviors, and suitabilities for targeted acquisition. Third, the pairwise framing allows the model to learn from within-protein comparisons in complex biological samples, where peptides from the same protein share the protein’s abundance, so observed intensity differences are attributable to peptide-specific response. A pairwise objective is also better matched to the assay-design decision than pointwise intensity regression: the user only needs to know which peptides are best within a given protein, not the absolute intensity any peptide will produce. This framing, combined with the use of complex biological samples, sidesteps the need for equimolar synthetic peptide datasets like those used to train PREGO ^20^.

Beyond these design choices, the training-size analysis indicates that Bromo can take advantage of additional data, whereas the XGBoost baseline quickly plateaus, suggesting that performance may continue to improve as larger and more diverse DIA resources become available. This scaling behavior also argues for prioritizing the curation of public DIA corpora as the most direct path to better peptide-ranking models.

The cross-instrument analyses emphasize that peptide response is not an intrinsic property of sequence alone. Replicate measurements acquired under the same instrument conditions yielded highly consistent peptide rankings, whereas rankings were substantially less consistent across different instrument platforms. This observation helps explain why a single universal peptide-ranking model is unlikely to be optimal for all experimental settings. Fine-tuning provides a practical way to adapt a pretrained model to new datasets, sample matrices, and instruments. In our experiments, fine-tuned models performed comparably to or better than models trained from scratch on most datasets, though the magnitude of improvement varied. This variation likely reflects two factors: the divergence between target and pretraining conditions, and the amount of representative data available for adaptation. In practice, Bromo should be viewed as a strong starting point for assay development; we recommend fine-tuning over training from scratch whenever DIA data from the target instrument are available, even at modest scale.

As with any supervised learning model, Bromo’s accuracy is limited by the size, quality, and diversity of its training data. The current model was trained and evaluated primarily using tryptic peptides and DIA-derived MS2 precursor-level response signals, so its performance may be reduced for other proteases, unusual peptide modifications, uncommon charge states, depletion or enrichment workflows, or sample matrices that are poorly represented in the training data. The model also inherits limitations from how labels are derived from DIA-NN output: missing peptide detections, interference, protein inference ambiguity, and intensity normalization can all affect the observed ranking used as ground truth. A related source of label noise arises when the assumption that peptides from the same protein share a common starting abundance is violated. A peptide assigned to a single protein or protein group may in fact be produced by more than one source, either by other isoforms of the same gene or by homologous proteins encoded by different genes that are not accurately represented in the protein database. In these cases, the peptide’s observed intensity includes contributions from all of these sources and no longer reflects the abundance of the protein to which it was assigned; endogenous post-translational modifications occurring at high stoichiometry act similarly, by depleting the unmodified form of the affected peptide in a manner that depends on cell state rather than on peptide sequence. Because these effects decouple a peptide’s effective starting abundance from the abundance of the protein it is assigned to, they are difficult to predict from the peptide sequence alone, and our pipeline mitigates them only partially, by excluding precursors that map to multiple proteins in the sequence database used and by aggregating labels across many runs. Even with high-quality data, replicate measurements on a single instrument yielded TKA values at *k* = 3 between 0.850 and 0.930, suggesting that the gold-standard ranking is itself noisy and that achievable model performance is bounded above by this replicate ceiling rather than by 1.0. Our comparison of binomial and majority-voting labeling schemes suggests that consensus labels are generally stable when multiple runs are available, but low-replicate datasets provide noisier supervision; this argues for prioritizing replicate depth over breadth when assembling future training corpora. More generally, future training sets that include a wider range of sample types, more instrument platforms, and carefully curated replicate measurements should improve both model accuracy and the reliability of fine-tuning.

Bromo solves only one part of targeted assay design. Selecting high-responding peptides is necessary, but a complete SRM or PRM method also requires choosing transitions or fragment ions, predicting or measuring retention times, avoiding interferences, and scheduling targets within instrument duty-cycle constraints. Practical deployment of Bromo should therefore combine its peptide-ranking output with tools for retention time prediction or calibration ^24,27–30^, fragment ion intensity prediction ^24,29–31^, transition selection, and scheduling optimization. A limitation of this study is that we did not use Bromo-selected peptides to develop and experimentally validate an SRM or PRM assay. Accordingly, our DIA-based evaluation establishes Bromo’s peptide-ranking performance, but not the performance of Bromo-selected peptides in a complete targeted workflow. Together, these components would allow Bromo to prioritize candidate peptides while downstream assay-design software determines which precursor and fragment ions can be measured reliably in a specific method.

Bromo is trained on DIA measurements acquired using fixed normalized collision energies and therefore does not model precursor-specific collision-energy optimization that may occur during targeted assay development. Because such optimization can alter fragmentation efficiency and the resulting quantitative signal, Bromo’s peptide rankings may not fully reflect the relative performance of candidate peptides in an optimized SRM or PRM method.

Overall, these results support the use of DIA data with deep proteome coverage and pairwise deep learning for peptide selection in targeted proteomics. Bromo improves sequence-based peptide ranking by aligning the learning objective with the assay-design decision, incorporating precursor charge state, and allowing adaptation to experiment-specific conditions. As public DIA datasets continue to grow and as peptide ranking is integrated with complementary prediction ^24,30^ and scheduling tools, Bromo should reduce the empirical burden of targeted assay development while preserving the flexibility needed for diverse biological and analytical applications.

## 4.1 Data availability

The paired human and yeast DIA MS/MS data with accession number PXD056793 ^24^ were downloaded from https://panoramaweb.org/Carafe.url. The large human cell line data was downloaded from PRIDE with accession number PXD030304. All search results, intermediate and processed data as well as model checkpoints used for reproducing the results in this paper are available through Zenodo with the link provided in the bromo-manuscript repository (https://github.com/Noble-Lab/bromo-manuscript).

## 4.2 Code availability

Bromo is open source and is available with an Apache 2.0 license at https://github.com/Noble-Lab/bromo. The scripts used to perform the analyses in this paper and reproduce figures and results are available at https://github.com/Noble-Lab/bromo-manuscript.

## Acknowledgments

This work was supported by the National Science Foundation award 2245300 (W.S.N.), the National Science Foundation Graduate Research Fellowship Program (Grant No. DGE-2140004, B.W.) and National Institutes of Health Grants R24 GM141156 and U01 DK137097 (M.J.M). During the preparation of this manuscript, the authors used Claude (Anthropic) and ChatGPT (OpenAI) to assist with language editing and improve the clarity of the text. The authors reviewed and edited all AI-assisted content and take full responsibility for the final manuscript.

## Author contributions

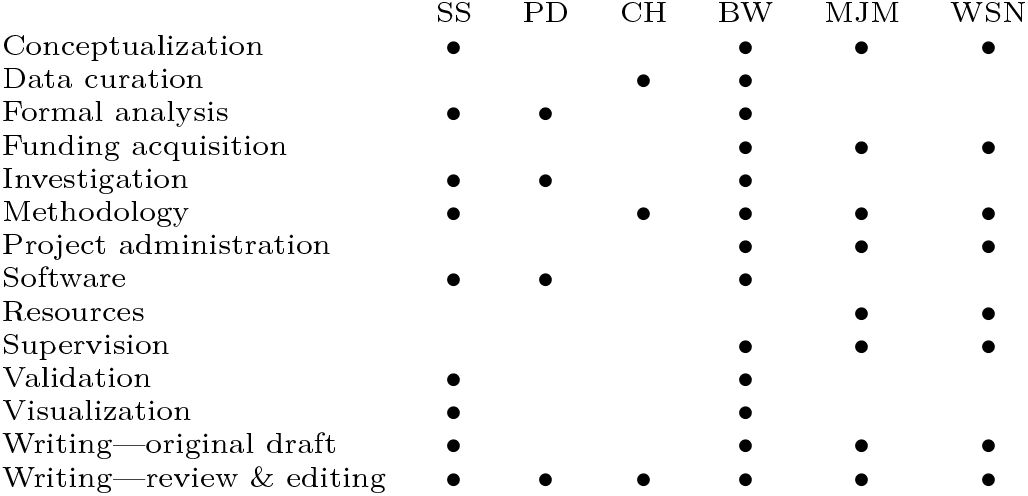

