## Supplemental Figures for "Prioritizing peptides for targeted mass spectrometry experiments using deep learning"

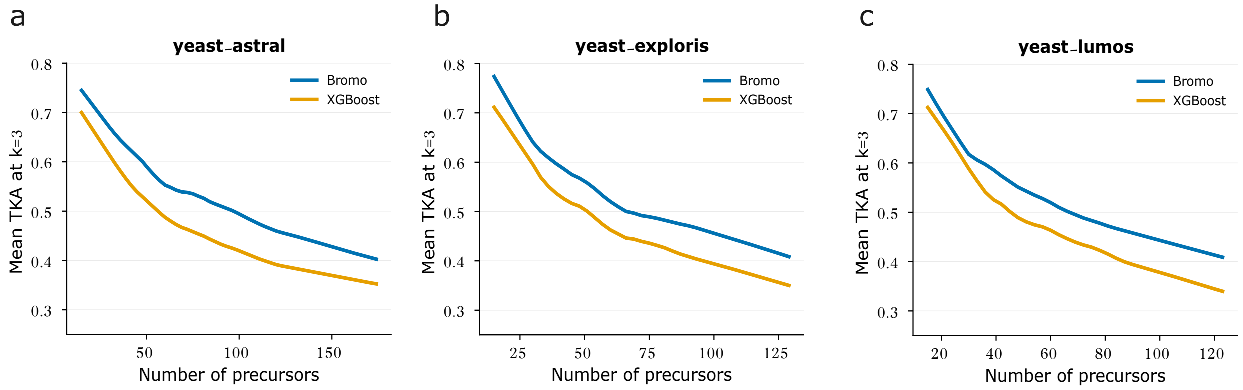


Supplementary Figure S1: **Peptide ranking performance on three DIA datasets as a function of number of precursors in protein group.** The figure plots the mean top-$k$ accuracy (TKA) for $k = 3$ as a function of number of precursors per protein group (Astral: **(a)**, Exploris: **(b)**, and Lumos: **(c)**) from Bromo (blue) and XGBoost (orange).

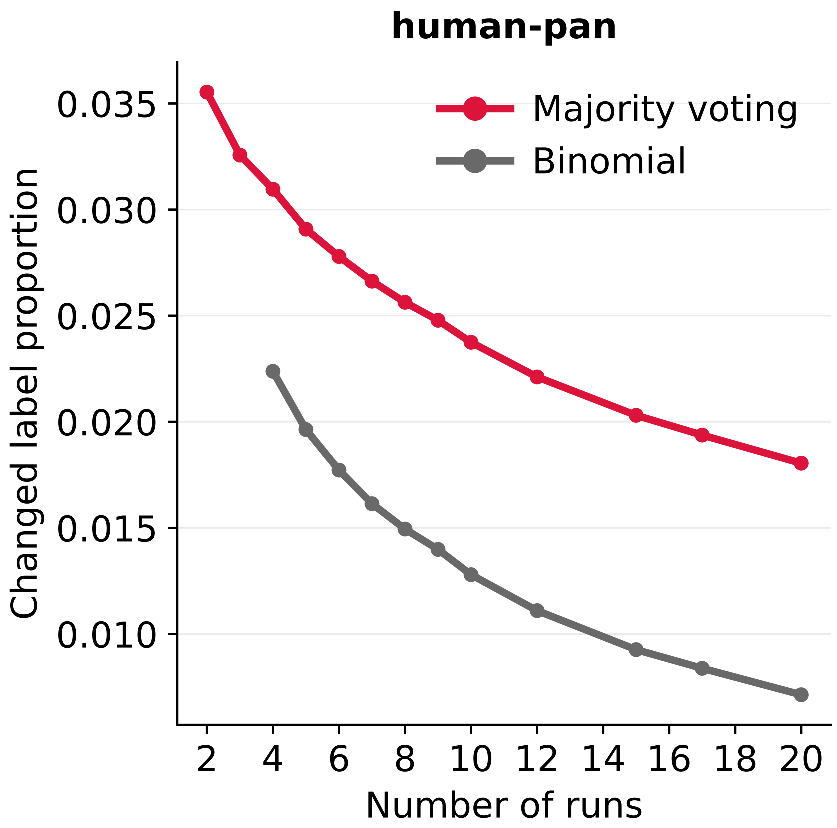


Supplementary Figure S2: **Comparing consistency of labeling schemes.** The figure plots the proportion of peptide pairs that receive different labels using the full human-pan dataset versus a downsampled version of the same data. The x-axis is the number of runs in the downsampled data, and the two series correspond to label assignment using the binomial distribution versus majority voting. With fewer than 5 runs, the binomial labeling scheme assigns all zero labels; thus, only the majority voting curve has values for fewer than 4 runs.

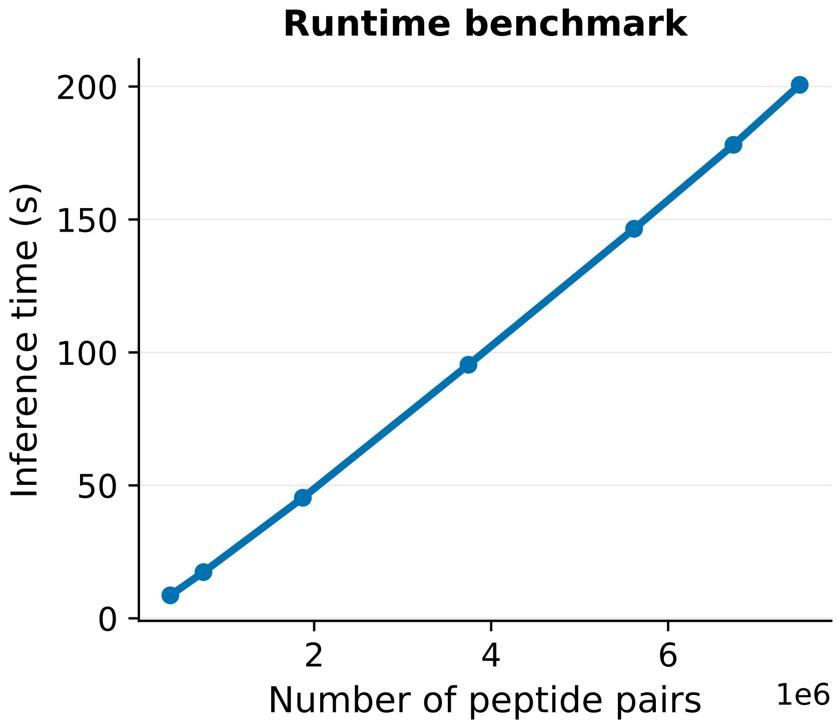


Supplementary Figure S3: **Inference runtime of Bromo across a range of subsampled dataset sizes.** Runtimes were measured on a single L40 GPU (48 GB) with 100 GB of CPU memory requested. The maximum GPU memory usage was 0.905 GB and maximum CPU memory usage was 0.253 GB.
